# Comprehensive Evaluation of Rapamycin’s Specificity as an mTOR Inhibitor

**DOI:** 10.1101/2022.12.10.519872

**Authors:** Filippo Artoni, Nina Grützmacher, Constantinos Demetriades

## Abstract

Rapamycin is a macrolide antibiotic that functions as an immunosuppressive and anti-cancer agent, and displays robust anti-ageing effects in multiple organisms including humans. Importantly, rapamycin analogs (rapalogs) are of clinical importance against certain cancer types and neurodevelopmental diseases. Although rapamycin is widely perceived as an allosteric inhibitor of mTOR (mechanistic target of rapamycin), the master regulator of cellular and organismal physiology, its specificity has not been thoroughly evaluated so far. In fact, previous studies in cells and in mice suggested that rapamycin may be also acting independently from mTOR to influence various cellular functions. Here, we generated a gene-edited cell line, that expresses a rapamycin-resistant mTOR mutant (mTOR^RR^), and assessed the effects of rapamycin treatment on the transcriptome and proteome of control or mTOR^RR^-expressing cells. Our data reveal a striking specificity of rapamycin towards mTOR, demonstrated by virtually no changes in mRNA or protein levels in rapamycin-treated mTOR^RR^ cells, even following prolonged drug treatment. Overall, this study provides the first comprehensive and conclusive assessment of rapamycin’s specificity, with important potential implications for ageing research and human therapeutics.

## Introduction

Rapamycin (also known as Sirolimus) is a naturally-occurring macrolide compound which was originally isolated from soil bacteria on Easter Island (Rapa Nui) in 1972 (Benjamin *et al*, 2011; Arriola Apelo & Lamming, 2016; Li *et al*, 2014). Rapamycin was primarily used in the clinic as an anti-fungal agent until 1999 when it was approved by the FDA for the prevention of kidney transplant rejection and later for the treatment of advanced kidney cell carcinoma. Its immunosuppressive and anti-proliferative properties are thought to be largely mediated by inhibition of mTOR, a serine/threonine kinase that functions as the master regulator of most cellular functions, including immune cell activation and cell growth control (Arriola Apelo & Lamming, 2016; Benjamin *et al*., 2011; Fernandes & Demetriades, 2021; Li *et al*., 2014). At the molecular level, rapamycin inhibits mTORC1 (mTOR complex 1) activity as a complex with the cytosolic immunophilin FKBP12 (FK506-binding protein 12). Binding of this drug-protein complex to mTOR blocks access to its catalytic site and prevents the phosphorylation of key substrates like S6K (ribosomal protein S6 kinase) (Chung *et al*, 1992). Additional immunophilins such as FKBP12.6, FKBP51, and FKBP52 have also been reported to bind and shape the pharmacology of rapamycin (Marz *et al*, 2013).

Over the last two decades, rapamycin has gained renewed interest after multiple studies uncovered its powerful anti-ageing properties (Arriola Apelo & Lamming, 2016; Benjamin *et al*., 2011; Fernandes & Demetriades, 2021; Li *et al*., 2014). Rapamycin has repeatedly been shown to extend lifespan and/or healthspan in worms (Robida-Stubbs *et al*, 2012), flies (Bjedov *et al*, 2010; Castillo-Quan *et al*, 2019; Schinaman *et al*, 2019), and mice (Bitto *et al*, 2016; Fok *et al*, 2014; Harrison *et al*, 2009). It extends chronological age in yeast (Powers *et al*, 2006) and reduces markers of senescence in cultured cells (Wang *et al*, 2017). More recently, studies conducted on marmoset monkeys have demonstrated rapamycin to have a good safety and tolerability profile (Lelegren *et al*, 2016; Tardif *et al*, 2015) thus paving the way for human trials. Currently, trials are underway on both companion dogs (Creevy *et al*, 2022) and humans (NCT04488601) with preliminary evidence suggesting improvement in cardiac function in middle-aged dogs who received rapamycin for 10 weeks (Urfer *et al*, 2017). Notably, rapamycin extends lifespan at doses much lower than the ones used to achieve immunosuppression in transplant patients thus minimizing potential side effects. Finally, even transient, intermittent, or late-life rapamycin administration has been shown to extend lifespan in flies and mice thus making Rapamycin an attractive option for human use as an anti-ageing compound (Bitto *et al*., 2016; Harrison *et al*., 2009; Partridge *et al*, 2020).

The target specificity of rapamycin has recently been debated. On one hand, *in vitro* kinase activity assays showed other serine/threonine kinases, besides mTOR, to be largely insensitive to rapamycin up to concentrations at the micromolar range. Likewise, receptor-binding assays indicated that many ligand-receptor interactions remain unaffected by rapamycin, with the possible exception of histamine I binding (European Medicine Agency, 2005). On the other hand, accumulating evidence in the literature suggests that rapamycin may exert some of its effects via mTOR-independent mechanisms. This is also supported by the fact that most kinase inhibitors demonstrate very low target selectivity (Hantschel, 2015; Hantschel *et al*, 2012; Karaman *et al*, 2008). For instance, rapamycin was suggested to block the exercise-induced accumulation of ribosomal RNA (rRNA) in the skeletal muscle of both wild-type and mice containing a rapamycin-resistant mTOR allele (Goodman *et al*, 2011). Moreover, rapamycin and other rapalogs were shown to directly bind and activate the lysosomal mucolipin TRP channel (TRPML1; also known as MCOLN1) independently of mTOR inhibition (Zhang *et al*, 2019). Finally, since FKBPs serve as chaperones for proper folding of several proteins (Bonner & Boulianne, 2017; Bultynck *et al*, 2001; Galfre *et al*, 2012; Vervliet *et al*, 2015; Wang *et al*, 1996), it can be speculated that their binding to rapamycin may be influencing cellular physiology via altering the interaction of FKBPs to their client proteins.

Here, to comprehensively address this important unresolved issue, we generated a rapamycin-resistant cell line by editing a single base in the *MTOR* gene at its endogenous locus (Choi *et al*, 1996; Hosoi *et al*, 1999; Lorenz & Heitman, 1995) and assessed how rapamycin affected the cellular transcriptome and proteome in an unbiased manner. These experiments unraveled an impressive specificity of rapamycin towards mTOR, with the rapamycin effects on gene expression and protein levels being virtually non-existent in the mTOR-mutant cells.

## Results

### Generation and characterization of a gene-edited cell line that expresses rapamycin-resistant mTOR

Mutations in the mTOR FRB (FKBP-rapamycin-binding) domain that disrupt its interaction with FKBP12 and confer rapamycin resistance to mTOR have been described almost 30 years ago (Brown *et al*, 1995; Chen *et al*, 1995; Choi *et al*., 1996; Lorenz & Heitman, 1995; Hara *et al*, 1997; Hosoi *et al*., 1999). Although such mTOR mutants have been used in previous studies, usually expressed exogenously in cells or via transgenic expression in mouse tissues, the presence of endogenous wild-type mTOR has complicated the interpretation of such results (Ge *et al*, 2009; Goodman *et al*., 2011; Luo *et al*, 2015; Zhang *et al*, 2000). Moreover, a comprehensive analysis of rapamycin’s effects in such cellular models, beyond single readouts, is lacking. Therefore, to probe whether rapamycin also acts through mTOR-independent mechanisms or exclusively via mTOR inhibition, we used CRISPR/Cas9-mediated gene editing to generate a HEK293FT cell line that expresses a Ser2035Thr mTOR mutant (Fig. 1A and Fig. EV1) that was previously described to be rapamycin-resistant (Brown *et al*., 1995; Chen *et al*., 1995; Choi *et al*., 1996; Lorenz & Heitman, 1995)(Hara *et al*., 1997; Hosoi *et al*., 1999). Intuitively, we called this mutant and the associated cell line, mTOR^RR^ (mTOR rapamycin-resistant). Importantly, as we mutated *MTOR* in the endogenous locus, no wild-type mTOR is expressed in these cells, verified by genomic DNA sequencing (Fig. EV1). Consistent with previous reports, the rapamycin-induced interaction of FLAG-tagged FKBP12 with wild-type mTOR, was completely abrogated in mTOR^RR^ (Fig. 1B). Of note, the catalytic activity of mTORC1 containing mTOR^RR^, as assessed by the phosphorylation of its direct substrate S6K, was indistinguishable from that in control cells and was diminished by treatment with Torin1, an ATP-competitive mTOR inhibitor (Fig. 1C). In contrast, mTOR^RR^ demonstrated complete resistance to rapamycin even when cells were treated with this compound at micromolar concentrations (Fig. 1C), or when the treatment was extended to 24 or 48 hours (Fig. 1D). Similar to rapamycin, mTOR^RR^ also exhibited full resistance to other rapalogs like everolimus and temsirolimus (Fig. 1E), but responded properly to amino acid (AA) starvation, showing that the regulation of mTORC1 by other inhibitory stimuli is unperturbed in mTOR^RR^ cells (Fig. 1F). In sum, the mTOR^RR^ HEK293FT cell line is a ‘clean’, reliable and robust model to investigate rapamycin’s specificity towards mTOR.

**Figure 1.**
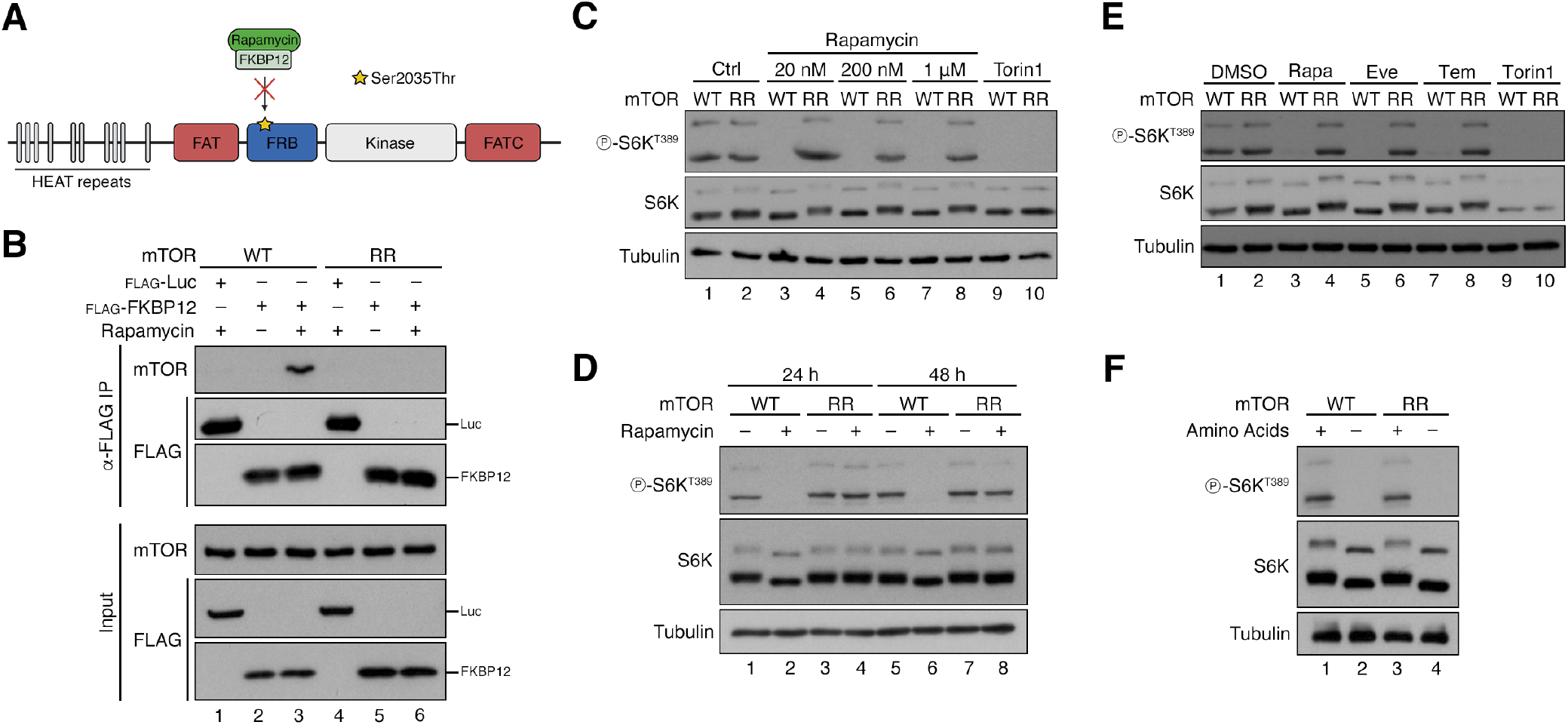
Generation and characterization of a gene-edited cell line that expresses rapamycin-resistant mTOR. **(A)** Schematic model of the mTOR protein depicting key domains and the location of the Ser2035Thr substitution in the FRB domain that prevents its inhibition by rapamycin/FKBP12. **(B)** Diminished binding of FKBP12 to the mTOR^RR^ mutant protein. Co-immunoprecipitation experiments in control (WT) or mTOR^RR^ HEK293FT cells, transiently expressing FLAG-tagged FKBP12 or Luciferase (Luc) as negative control, treated with rapamycin (20 nM, 1 h) or DMSO. Binding of mTOR to FKBP12 was analyzed by immunoblotting as indicated. **(C)** mTOR^RR^ does not respond to rapamycin even at extremely high concentrations. Immunoblots with lysates from HEK293FT WT and mTOR^RR^ cells treated for 1 h with 20 nM to 1 μM rapamycin as indicated, 250 nM Torin1, or DMSO as control. mTORC1 activity was assessed by S6K phosphorylation. **(D)** mTOR^RR^ is resistant to long-term rapamycin treatment. Immunoblots with lysates from HEK293FT WT and mTOR^RR^ cells treated with DMSO or rapamycin (20 nM) for 24 or 48 h, probed with the indicated antibodies. **(E)** mTOR^RR^ shows resistance to multiple rapalogs. Immunoblots with lysates from HEK293FT WT and mTOR^RR^ cells treated with DMSO, rapamycin (Rapa; 20 nM), everolimus (Eve; 20 nM), temsirolimus (Tem; 20 nM), or Torin1 (250 nM) for 1 h, probed with the indicated antibodies. **(F)** Cell expressing mTOR^RR^ have active mTORC1 that responds properly to AA starvation. Immunoblots with lysates from HEK293FT WT and mTOR^RR^ cells treated with control (+) or AA-free media (−) for 1 h, probed with the indicated antibodies.

### Rapamycin alters gene expression exclusively via mTOR inhibition

Having validated our experimental model, we then treated control (WT) and mTOR^RR^ cells with rapamycin for 24 hours and performed RNA-seq experiments to investigate its effects on global gene expression. In line with mTOR—directly or indirectly—regulating the activity of several transcription factors (Hardwick *et al*, 1999; Laplante & Sabatini, 2013), we detected more than 5000 genes whose expression changed significantly in WT cells upon rapamycin treatment (Fig. 2A,B and Suppl. Table 1). Our analysis identified several genes that are known to be affected by mTOR inhibition (e.g., HMOX1, RHOB, MYC) (Bayeva *et al*, 2012; Gordon *et al*, 2015; Jin *et al*, 2013; Sun *et al*, 2022; Visner *et al*, 2003) (Fig. 2B) thus validating our experimental setup. Similarly, rapamycin decreased the expression of AA transporters like SLC7A5 and SLC7A11, which are known to be regulated downstream of an mTORC1-ATF4 axis (Torrence *et al*, 2021) (Fig. 2B and Suppl. Tables 1, 2).

**Figure 2.**
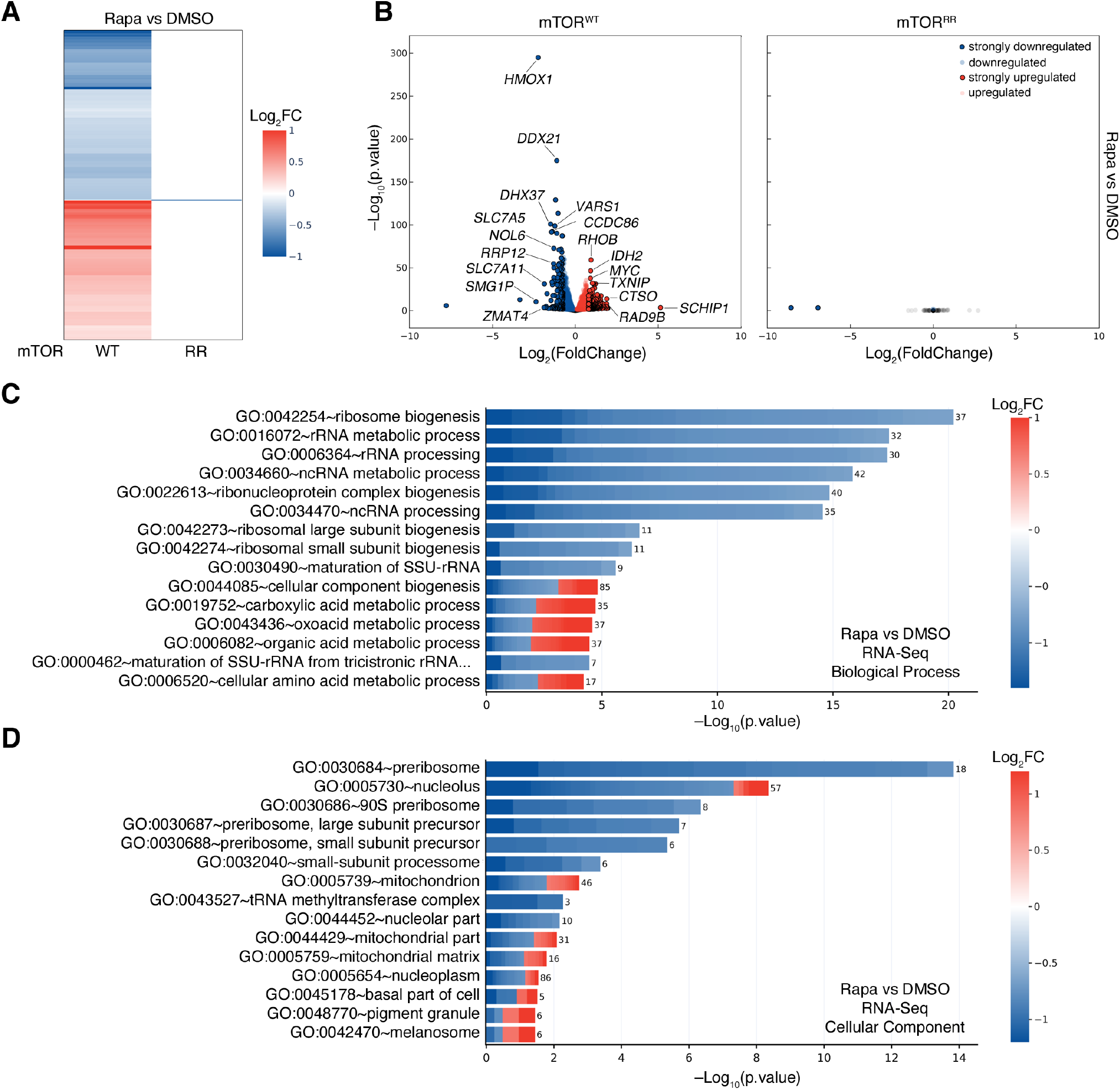
Rapamycin alters gene expression exclusively via mTOR inhibition. **(A)** Heatmap depicting the rapamycin-induced changes in gene expression in HEK293FT WT and mTOR^RR^ cells. RNA-Seq data from cells treated with rapamycin (20 nM, 24 h) expressed as log-transformed fold change (Log_2_FC). Only statistically-significant changes (p < 0.05) are shown. **(B)** Volcano plots showing the rapamycin-induced changes in gene expression in HEK293FT WT (left) and mTOR^RR^ (right) cells from the RNA-Seq experiment described in (A). Genes that are significantly (p < 0.05) down- or upregulated by rapamycin are shown in blue or red respectively. Strongly down- (Log_2_FC < −0.75) or upregulated (Log_2_FC > +0.8) genes are shown with black outline. Unaffected genes (p > 0.05) shown as grey dots. Selected genes are marked in the plot. **(C)** Biological process (BP) GO term analysis using the genes that are strongly down- (blue) or upregulated (red) by rapamycin in WT cells as described in (B). The color of each box in the cell plot represents log-transformed fold change values for each gene in rapamycin- vs DMSO-treated cells. The number of genes in the selected dataset for each GO term is shown on the right side of each bar. **(D)** As in (C), but for Cellular Component (CC) GO term analysis.

Gene ontology (GO) term enrichment analysis, using the differentially regulated genes that are strongly down- or upregulated by rapamycin (Log_2_FC < –0.75 and Log_2_FC > +0.8, respectively), revealed a strong enrichment of ribosome-related Biological Process (BP) and Cellular Component (CC) terms (e.g., BP:GO:0042254~ribosome biogenesis; CC:GO:0030684~preribosome) (Fig. 2C,D and Suppl. Table 2). Confirming the well-known role of mTOR in promoting ribosome biogenesis and positively regulating rRNA expression (Mayer & Grummt, 2006; Powers & Walter, 1999), all genes that fall under these terms (e.g., RRP12, RRP9, RRS1, RRP1, RPF2, NOP16, NOL6, DDX21, MRTO4, MRM1) were found to be downregulated by rapamycin (Fig. 2C,D and Suppl. Table 2). Interestingly, we also observed a strong enrichment of terms related to mitochondria-resident proteins (e.g., CC:GO:0005739~mitochondrion; CC:GO:0005759~mitochondrial matrix) and associated mitochondrial functions (e.g., BP:GO:0019752~carboxylic acid metabolic process) among the genes that are differentially regulated by rapamycin (Fig. 2C,D and Suppl. Table 2). Similar results were obtained when performing the GO analysis with more relaxed criteria including all significantly down- and upregulated genes, instead of setting cut-offs for those that change robustly upon rapamycin (Fig. EV2A,B and Suppl. Table 3). Strikingly, unlike the massive transcriptional effects of rapamycin in control cells, we found only 3 genes whose expression was altered in mTOR^RR^ cells treated with rapamycin (Fig. 2A,B and Suppl. Table 1). These data indicated that rapamycin practically regulates transcription exclusively via its direct inhibitory effect on mTOR.

### Rapamycin alters the cellular proteome exclusively through mTOR

In addition to their involvement in transcriptional regulation, the best-described role of rapamycin and mTORC1 is in the control of *de novo* protein synthesis via controlling the phosphorylation and activity of—direct or indirect—mTORC1 targets like S6K, 4E-BP1, and S6, with only certain 4E-BP1 phospho-sites being rapamycin-sensitive (Thoreen *et al*, 2012; Thoreen *et al*, 2009). Therefore, we next sought to investigate the rapamycin-induced changes in the cellular proteome and to explore how much of this happens due to mTOR inhibition.

To this end, we treated control and mTOR^RR^ cells with rapamycin for 24 or 48 hours and performed whole-proteome quantitative mass spectrometry experiments. Out of a total of approximately 7500 proteins that were detected and quantified, the levels of more than 2500 and 2300 proteins changed significantly in WT cells treated with rapamycin for 24 or 48 hours, respectively (Fig. 3A-C and Suppl. Table 4). Similar to what we observed in transcriptomic analyses, these hits include multiple proteins belonging to pathways that have been previously described to be regulated by rapamycin and mTOR (e.g., PDCD4, RHOB, HMOX1, SQSTM1, PRELID1, SCD, SESN2) (Bayeva *et al*., 2012; Dorrello *et al*, 2006; Gordon *et al*., 2015; Jin *et al*., 2013; Ko *et al*, 2017; Mauvoisin *et al*, 2007; Sun *et al*., 2022; Visner *et al*., 2003; Wall *et al*, 2008; Zhu *et al*, 2020) as well as several amino acid transporters (SLC7A11, SLC38A10, SLC3A2, SLC7A5) (Graber *et al*, 2017; Nachef *et al*, 2021; Torrence *et al*., 2021; Zhang *et al*, 2021) (Fig. 3B,C and Suppl. Table 4). Remarkably, however, none of the 7574 detected proteins were differentially regulated in rapamycin-treated mTOR^RR^ cells, even after prolonged drug treatment (Fig. 3A-C and Suppl. Table 4), again showing complete absence of mTOR-independent effects by rapamycin.

**Figure 3.**
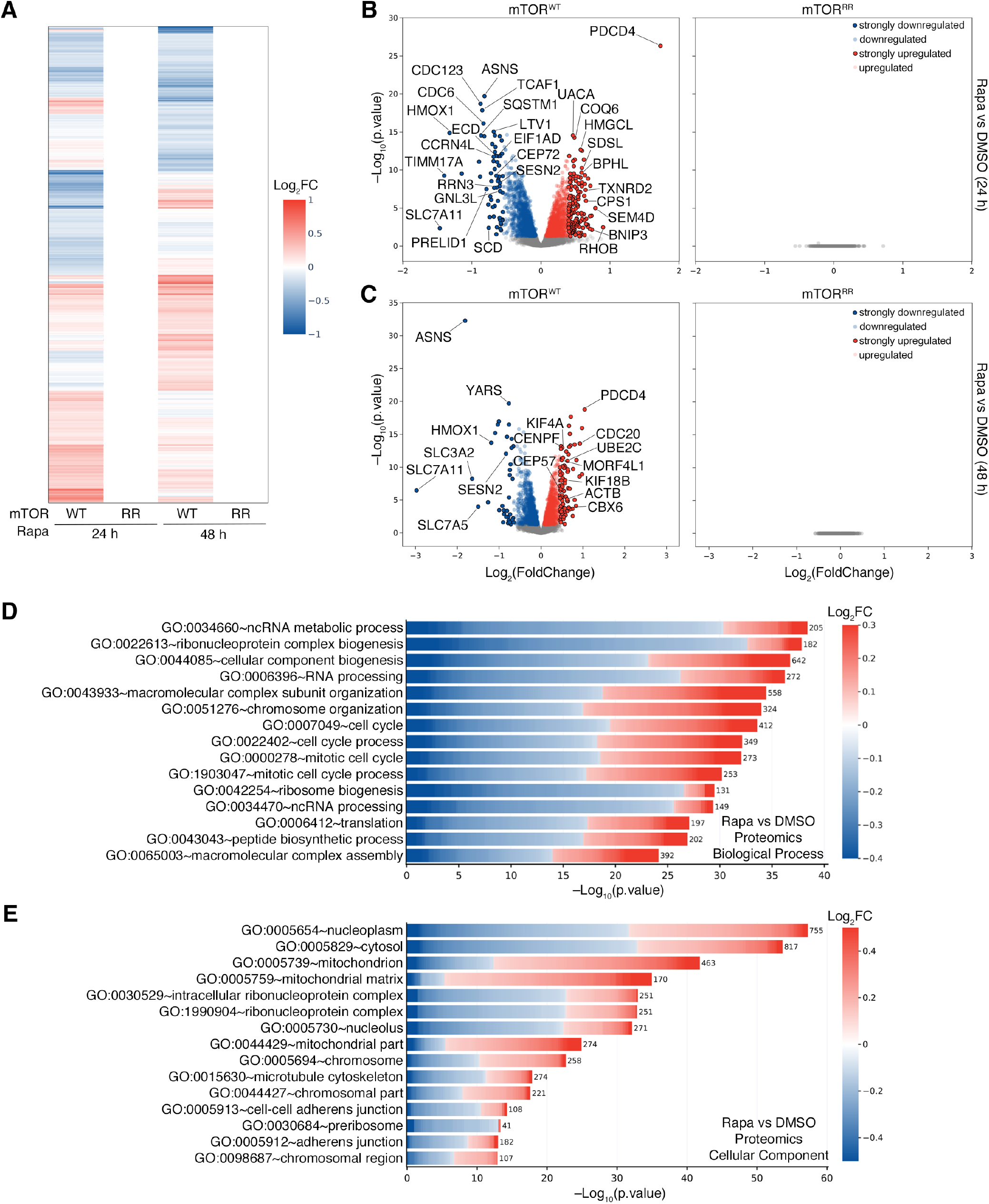
Rapamycin alters the cellular proteome exclusively via mTOR inhibition. **(A)** Heatmap depicting the rapamycin-induced changes in the proteome of HEK293FT WT and mTOR^RR^ cells. Shown are whole-proteome quantitative mass-spectrometry data from cells treated with rapamycin (20 nM) for 24 or 48 h expressed as log-transformed fold change (Log_2_FC). Only statistically-significant changes (p < 0.05) are shown. **(B)** Volcano plots showing the rapamycin-induced proteomic changes in HEK293FT WT (left) and mTOR^RR^ (right) cells from the mass-spectrometry experiment described in (A). Proteins that are significantly (p < 0.05) down- or upregulated by rapamycin (20 nM, 24 h) are shown in blue or red respectively. Most strongly down- (Log_2_FC < –0.55) or upregulated (Log_2_FC > +0.4) proteins are shown with black outline. Unaffected proteins (p > 0.05) shown as grey dots. Selected proteins are marked in the plot. **(C)** As in (B), but for cells treated with 20 nM rapamycin (or DMSO) for 48 h. Strongly down- (Log_2_FC < –0.65) or upregulated (Log_2_FC > +0.45) proteins are shown with black outline. **(D)** Biological process (BP) GO term analysis using the proteins that are significantly down- (blue) or upregulated (red) by 24 h rapamycin in WT cells as described in (B). The color of each box in the cell plot represents log-transformed fold change values for each protein in rapamycin- vs DMSO-treated cells. The number of proteins in the selected dataset for each GO term is shown on the right side of each bar. **(E)** As in (D), but for Cellular Component (CC) GO term analysis.

Consistent with our observations from RNA-Seq experiments, GO analysis using all proteins that are significantly down- or upregulated (p < 0.05) upon 24-hour rapamycin treatment in WT cells showed strong enrichment of terms related to ribosomes and translation (e.g., BP:GO:0022613~ribonucleoprotein complex biogenesis; BP:GO:0042254~ribosome biogenesis; BP:GO:0006412~translation) (Fig. 3D and Suppl. Table 5) with the majority of proteins that belong to this category being downregulated (e.g., CDC123, CCRN4L, LTV1, RRN3, EIF1AD, SESN2, GNL3L, CCDC86, WDR74, TNRC6A) (Fig. 3B,D and Suppl. Tables 4, 5). Another prominent group of GO terms that is enriched in the proteomic analysis includes terms related to the cell cycle (e.g., BP:GO:0007049~cell cycle, BP:GO:0000278~mitotic cell cycle) (Fig. 3D and Suppl. Tables 4, 5) with proteins being either down- or upregulated upon rapamycin treatment (e.g., CDC123, CDC6, ASNS, ECD, CEP72, PDCD4, RHOB, HMGN3, DHFR, TRIOBP) (Fig. 3D). Of note, this is in line with the well-established role of rapamycin and mTOR in the regulation of cell proliferation (Dowling *et al*, 2010). Similar results were obtained when analyzing the respective dataset (all significantly-regulated proteins; p < 0.05) from 48-hour rapamycin treatment in WT cells (Fig. 3A,C, Fig. EV3 and Suppl. Table 6).

Furthermore, in line with the robust expression changes in genes related to mitochondria (Fig. 2C,D), we observed similar effects in the levels of mitochondrial proteins, as shown by the strong enrichment of related GO terms (e.g., CC:GO:0005739~mitochondrion; CC:GO:0005759~mitochondrial matrix) (Fig. 3E and Suppl. Tables 4, 5). Interestingly, when performing the GO term analysis using only the strongly up- or downregulated proteins, the enrichment of terms related to mitochondrial proteins became even more prominent (Fig. EV4 and Suppl. Table 7) with most mitochondria-related proteins being upregulated by rapamycin (e.g., BNIP3, BNIP3L, CPS1, TXNRD2, SDSL, ACSF2, BPHL, HMGCL, HSD17B8, SFN) (Fig. EV4 and Suppl. Tables 4, 7).

Finally, although a connection between mTOR activity and cell adhesion has been described before, the underlying mechanisms are less clear (Asrani *et al*, 2017; Chen *et al*, 2015). Interestingly, our proteomics analysis showed that rapamycin treatment led to significant changes in the levels of a large number of proteins that are related to adherens and anchoring junctions (CC:GO:0005912~adherens junction; CC:GO:0070161~anchoring junction), most of which are downregulated under these conditions (e.g., GJA1, SLC3A2, RANGAP1, DSP, FASN, APC, EEF2, EIF2A, TNKS1BP1, ATXN2L) (Fig. 3B,E and Suppl. Tables 4, 7). This suggests that some of the effects of rapamycin/mTOR on cell adhesion may stem from changes in the junctional proteome.

In sum, our unbiased interrogation of rapamycin’s effects in cells reveals an extraordinary specificity of this compound towards mTOR, and—at the same time—provides a comprehensive evaluation of its role on the cellular transcriptome and proteome.

## Discussion

According to previous studies, the majority of inhibitory compounds that are used in research or in therapeutics demonstrate off-target effects, influencing the activity of additional signaling molecules that can also be structurally-unrelated to their presumed targets. Sometimes, inhibitors even show higher potency towards off-targets compared to on-target effects (Davies *et al*, 2000; Bain *et al*, 2003; Bain *et al*, 2007; Fedorov *et al*, 2007; Davis *et al*, 2011). In fact, very few FDA-approved kinase inhibitors have demonstrated high target selectivity, whereas the majority influences the activity of 10-100 off-target kinases (Hantschel, 2015; Hantschel *et al*, 2012; Karaman *et al*, 2008). Importantly, the low specificity and selectivity of most kinase inhibitors greatly complicates the interpretation of data originating from their use and has important implications for their applicability in both research and therapeutics.

Surprisingly, although rapamycin has been used as an mTOR inhibitor for more than three decades, its specificity towards this key signaling hub has not been thoroughly evaluated so far. In fact, previous studies have hinted at the existence of mTOR-independent functions of rapamycin in cells and in transgenic mouse models thus underscoring the need for a detailed evaluation of its specificity. In transgenic mice overexpressing a rapamycin-resistant mTOR mutant in skeletal muscles, rapamycin was still able to partially suppress mechanical-loading-induced ribosome biogenesis (Goodman *et al*., 2011). It is worth noting however, that these transgenic mice were maintained as hemizygotes with the mTOR^RR^ allele expressed on top of endogenous wild-type mTOR (Ge *et al*., 2009; Goodman *et al*., 2011), which does not allow for safe conclusions to be drawn from such experiments. Indeed, the observed rapamycin effects that were previously interpreted as mTOR-independent can also be ascribed to the inhibition of endogenous wild-type mTOR molecules. This is also supported by our findings, using a gene-edited cell line that expresses mutant, rapamycin-resistant mTOR from the endogenous *MTOR* locus, while lacking expression of wild-type mTOR: unlike the previously-described mTOR^RR^ transgenic mice, our mTOR^RR^ cells are fully resistant to rapamycin, which is demonstrated by unaffected mTORC1 activity, diminished FKBP12-mTOR binding, and complete lack of transcriptional or proteomic changes in response to rapamycin treatment.

A more recent study reported that micromolar concentrations of rapamycin (or of other rapalogs like everolimus and temsirolimus) can directly bind to TRPML1/MCOLN1, the primary lysosomal calcium release channel, and enhance its activity in an mTOR-independent manner (Zhang *et al*., 2019). Although rapalogs may indeed act on additional substrates at such concentrations, these are two to three orders of magnitude higher than those classically used in cell culture experiments (2-20 nM; like those we used for the transcriptomic and proteomic analyses described here) or—most importantly—those that are detected in the blood of patients that take rapamycin. More specifically, whole-blood concentrations (WBC) of rapamycin/sirolimus are in the 10-20 nM range in renal transplant patients to prevent graft rejection (Meier-Kriesche & Kaplan, 2000). In patients treated with everolimus against metastatic renal cell carcinoma, the median WBC is approximately 15 nM (Takasaki *et al*, 2019), while plasma concentrations of temsirolimus for relapsed/refractory multiple myeloma average around 9 nM (Farag *et al*, 2009). Higher doses of temsirolimus have only been used as a last resort in clinical trials against aggressive forms of lymphoma with WBC values reaching 500-600 nM (Hudes *et al*, 2007), which is still more than 25 times lower than the rapamycin concentration required to half-maximally activate TRPML1 (Zhang *et al*., 2019). Thus, the rapamycin-mediated TRPML1 activation, while observed *in vitro*, is not likely to be physiologically relevant for research and therapeutics. Furthermore, rapamycin doses aimed at slowing or reversing ageing are generally even lower than those used for immunosuppression and cancer treatment (Bitto *et al*., 2016; Bjedov *et al*., 2010; Castillo-Quan *et al*., 2019; Fok *et al*., 2014; Harrison *et al*., 2009; Robida-Stubbs *et al*., 2012; Schinaman *et al*., 2019), further suggesting that TRPML1 activation is unlikely to be involved in rapamycin-mediated lifespan extension.

Rapamycin acts an allosteric mTOR inhibitor in a complex with FKBP12 and other FKBP family members (Chung *et al*., 1992; Marz *et al*., 2013). Most FKBPs possess peptidyl-prolyl isomerase (PPIase) activity, hence functioning as protein folding chaperones for a variety of different proteins (Harrar *et al*, 2001; Kolos *et al*, 2018). For instance, FKBP12 and FKBP12.6 have been shown to bind and modulate the activity of ryanodine receptors (RyRs) and inositol 1,4,5-trisphosphate receptors (IP3Rs) (Bultynck *et al*., 2001; Galfre *et al*., 2012; Vervliet *et al*., 2015), which are channels involved in intracellular calcium release. FKBP12 has also been shown to inhibit TGFβ family type I receptors (Wang *et al*., 1996). Likewise, the larger FKBP51 and FKBP52 chaperones control glucocorticoid receptor (GR) localization and activity (Fries *et al*, 2017) as well as the protein levels of Argonaute 2 (AGO2) (Martinez *et al*, 2013), an essential component of the RNA-induced silencing complex (RISC). Hence, it would be reasonable to speculate that binding of rapamycin to different FKBP proteins may influence the folding—and thus function—of client proteins, also beyond mTOR inhibition. Because any changes in receptor or signaling pathway activities, organelle function, metabolism, or other cellular processes are eventually translated to changes in gene or protein expression, we here analyzed the cellular transcriptome and proteome to interrogate the effects of rapamycin treatment in an unbiased manner. These experiments revealed a striking dependence of rapamycin on mTOR inhibition to influence cells, with mTOR^RR^-expressing cells demonstrating virtually no effects upon treatment with this drug. In sum, although rapamycin could—in theory—affect cells also independently from mTOR inhibition (via TRPML1 or through FKBP-dependent mechanisms), this is not the case, at least for rapamycin concentrations that are within the nanomolar range used in research or found in the blood of patients treated with this drug or its analogs.

In addition to assessing rapamycin’s specificity towards mTOR, we here also investigated how rapamycin influences gene expression and whole-cell protein levels in control cells that express wild-type mTOR. Consistent with the role of mTORC1 in regulating cap-dependent translation (Fonseca *et al*, 2014; Holz *et al*, 2005; Liu & Sabatini, 2020; Ma & Blenis, 2009; Thoreen *et al*., 2012) and with rapamycin’s ability to repress both global and specific translation of key subsets of transcripts (Dickinson *et al*, 2011; Huo *et al*, 2011; Nandagopal & Roux, 2015; Tsukumo *et al*, 2016; Wang *et al*, 2007), these transcriptomics and proteomics datasets identified several thousands of genes and proteins whose expression changes in rapamycin-treated cells. For instance, highlighting the well-described role of rapamycin/mTOR on ribosomal biogenesis, rapamycin treatment strongly downregulated the expression of genes encoding for ribosomal proteins, tRNA synthetases (Kim *et al*, 2017; Lee *et al*, 2012) and other accessory proteins involved in this process (Mayer & Grummt, 2006; Powers & Walter, 1999) (Fig. 2 and EV2). Accordingly, we observed a strong enrichment of proteins related to non-coding RNA metabolic processes and ribonucleoprotein complex biogenesis among those downregulated by rapamycin (Fig. 3 and EV3). Furthermore, both our RNA-seq and proteomics data also confirm previous studies about rapamycin’s role in regulating mitochondrial function (Morita *et al*, 2017; Ramanathan & Schreiber, 2009; Rosario *et al*, 2019; Schieke *et al*, 2006; Villa-Cuesta *et al*, 2014) (Fig. 2, 3 and EV2, 3). Interestingly, we find that proteins that are associated with adherens/anchoring junctions are enriched among those downregulated by rapamycin, which may suggest a role for mTOR in the regulation of cell-cell and cell-matrix contacts (Fig. 3E and EV3B). Finally, we observe an enrichment of terms associated with cell cycle and mitosis in the rapamycin-dependent proteome, possibly reflecting the anti-proliferative effects of rapamycin (Zaragoza *et al*, 1998) and the role of mTOR in cell proliferation (Dowling *et al*., 2010).

Overall, we here provide an unbiased, comprehensive evaluation of rapamycin’s specificity towards mTOR in mammalian cells, in unprecedented depth. Given the immense potential that rapamycin and its analogs have as anti-ageing compounds or in therapeutics, these findings provide important insight for both basic and translational research and aim at improving the applicability and specificity of its use in humans.

## Methods

### Cell culture

All cell lines were grown at 37 ^o^C, 5% CO2. Human female embryonic kidney HEK293FT cells (#R70007, Invitrogen; RRID: CVCL_6911) were cultured in high-glucose Dulbecco’s Modified Eagle Medium (DMEM) (#41965-039, Gibco) supplemented with 10% fetal bovine serum (FBS) (#F7524, Sigma, or #S1810, Biowest) and 1% Penicillin-Streptomycin (#15140-122, Gibco).

HEK293FT cells were purchased from Invitrogen. The identity of the HEK293FT cells was validated by the Multiplex human Cell Line Authentication test (Multiplexion GmbH), which uses a single nucleotide polymorphism (SNP) typing approach, and was performed as described at www.multiplexion.de. All parental and edited cell lines were regularly tested for *Mycoplasma* contamination using a PCR-based approach and were confirmed to be *Mycoplasma-free*.

### Cell culture treatments

To allosterically inhibit mTOR, rapamycin (#S1039, Selleckchem), everolimus (#S1120, Selleckchem) or temsirolimus (# S1044, Selleckchem) were dissolved in DMSO and added directly into full cell culture media at a final concentration of 20 nM unless otherwise indicated in the figure legends. Treatments were performed for the times described in the figure legends. DMSO was used as a negative control for all treatments. Torin1 (#4247, Tocris) was used as an ATP-competitive mTOR inhibitor and added in the culture media at a final concentration of 250 nM for 1 hour.

Amino acid (AA) starvation experiments were performed as described previously (Demetriades *et al*, 2014; Demetriades *et al*, 2016). In brief, custom-made starvation media were formulated according to the Gibco recipe for high-glucose DMEM specifically omitting all AAs. The media were filtered through a 0.22-μm filter device and tested for proper pH and osmolality before use. For the respective AA-replete (+AA) treatment media, commercially available high-glucose DMEM was used (#41965039, Thermo Fisher Scientific). All treatment media were supplemented with 10% dialyzed FBS (dFBS) and 1x Penicillin-Streptomycin (#15140-122, Gibco). For this purpose, FBS was dialyzed against 1x PBS through 3,500 MWCO dialysis tubing. For basal (+AA) conditions, the culture media were replaced with +AA treatment media 1 hour before lysis. For amino-acid starvation (−AA), culture media were replaced with starvation media for 1 hour.

### Antibodies

Antibodies against phospho-p70 S6K (Thr389) (#9205), p70 S6K (#9202 for Fig. 1C,E; #97596 for Fig. 1D,F), FLAG (#2368) and mTOR (#2983) were purchased from Cell Signaling Technology. The anti-human tubulin (#T9026) antibody was purchased from Sigma.

### Plasmid DNA transfections

Plasmid DNA transfections in HEK293FT cells were performed using Effectene transfection reagent (#301425, QIAGEN) according to the manufacturer’s instructions.

### Generation of the gene-edited mTOR^RR^ cell line

The rapamycin-resistant mTOR (mTOR^RR^) HEK293FT cell line was generated by gene-editing using the pX459-based CRISPR/Cas9 method as described elsewhere (Ran *et al*, 2013). The sgRNA expression vectors were generated by cloning appropriate DNA oligonucleotides (Suppl. Table 8) in the BbsI restriction sites of pX459 (#62988, Addgene). In brief, transfected cells were selected with 3 μg/ml puromycin (#A11138-03, Thermo Fisher Scientific) 48 hours post-transfection. Single-cell clones were generated by FACS sorting and individual clones were validated by genomic DNA sequencing and functional assays.

### Gene expression analysis (RNA-Seq)

To analyze gene expression changes via RNA-seq experiments, total mRNA was isolated using QIAshredder columns (#79656, QIAGEN) and the RNeasy Plus Mini Kit (#74034, QIAGEN) according to the manufacturer’s instructions. RNA-seq experiments were performed by the Max Planck Genome Centre (MPGC) Cologne, Germany (https://mpgc.mpipz.mpg.de/home/). RNA quality was assessed with an Agilent Bioanalyzer (Nanochip). Library preparation was done according to NEBNext Ultra™ II Directional RNA Library Prep Kit for Illumina (#E7760L, New England Biolabs) including polyA enrichment and addition the of ERCC RNA spike-ins. Libraries were quality controlled by Agilent TapeStation or LabChip GX or GX Touch (PerkinElmer). Sequencing-by-synthesis was performed on a HiSeq 3000 (Illumina) with single read mode 1 x 150 bp. Data from one representative RNA-seq experiment, out of two independent replicates, are shown in this manuscript. Each experiment was performed from 3 independent biological replicates. The raw data from both RNA-seq experiments are available in the NCBI Sequence Read Archive (see also the Data Availability section).

### Cell lysis and immunoblotting

For standard SDS-PAGE and immunoblotting experiments, cells from a well of a 12-well plate were treated as indicated in the figures and lysed in 250 μl of ice-cold Triton lysis buffer (50 mM Tris pH 7.5, 1% Triton X-100, 150 mM NaCl, 50 mM NaF, 2 mM Na-vanadate, 0.011 gr/ml beta-glycerophosphate) supplemented with 1x PhosSTOP phosphatase inhibitors (#4906837001, Roche) and 1x cOmplete protease inhibitors (#11836153001, Roche) for 10 minutes on ice. Samples were clarified by centrifugation (15000 rpm, 10 min, 4 °C) and supernatants were boiled in 1x SDS sample buffer (5x SDS sample buffer: 350 mM Tris-HCl pH 6.8, 30% glycerol, 600 mM DTT, 12.8% SDS, 0.12% bromophenol blue). Protein samples were subjected to electrophoretic separation on SDS-PAGE and analyzed by standard Western blotting techniques. In brief, proteins were transferred to nitrocellulose membranes (#10600002 or #10600001, Amersham) and stained with 0.2% Ponceau solution (#33427-01, Serva) to confirm equal loading. Membranes were blocked with 5% skim milk powder (#42590, Serva) in PBS-T [1x PBS, 0.1% Tween-20 (#A1389, AppliChem)] for 1 hour at room temperature, washed three times for 10 min with PBS-T and incubated with primary antibodies [1:1000 in PBS-T, 5% bovine serum albumin (BSA; #10735086001, Roche)] rotating overnight at 4 °C. The next day, membranes were washed three times for 10 min with PBS-T and incubated with appropriate HRP-conjugated secondary antibodies (1:10000 in PBS-T, 5% milk) for 1 hour at room temperature. Signals were detected by enhanced chemiluminescence (ECL) using the ECL Western Blotting Substrate (#W1015, Promega) or SuperSignal West Pico PLUS (#34577, Thermo Scientific) and SuperSignal West Femto Substrate (#34095, Thermo Scientific) for weaker signals. Immunoblot images were captured on films (#28906835, GE Healthcare; #4741019289, Fujifilm).

### Co-immunoprecipitation (co-IP)

For co-IP experiments, 1.5 x 10^6^ cells were transiently transfected with the indicated plasmids and lysed 36 h post-transfection in IP lysis buffer (50 mM Tris pH 7.5, 0.3% CHAPS, 150 mM NaCl, 50 mM NaF, 2 mM Na-vanadate, 0.011 g/ml β-glycerophosphate, 1x PhosSTOP phosphatase inhibitors and 1x cOmplete protease inhibitors). FLAG-tagged proteins were incubated with 30 μl pre-washed anti-FLAG M2 affinity gel (Sigma, #A2220) for 3 h at 4 °C and washed four times with IP wash buffer (50 mM Tris pH 7.5, 0.3% CHAPS, 150 mM NaCl and 50 mM NaF). Samples were then boiled for 6 min in 1x SDS sample buffer and analysed by immunoblotting using appropriate antibodies.

### Quantitative whole-cell proteomics

For mass-spectrometry experiments, HEK293FT cells were cultured in 10 cm dishes in 10 ml of complete culture media as described above. Rapamycin (or DMSO as negative control) were added directly in the culture media for 24 or 48 hours. Experiments were performed with 5 independent biological replicates per condition (1x 10 cm dish per replicate). In brief, cells were scraped, collected in 1.5 ml tubes on ice, washed in serum-free media, pelleted by centrifugation (500 x g, 3 min), and cell pellets were snap-frozen in liquid nitrogen and stored at −80 °C.

#### Sample preparation

For sample preparation for quantitative proteomic analysis, cell pellets were lysed in 6 M guanidinium chloride (GdmCl) supplemented with 2.5 mM TCEP (tris(2-carboxyethyl)phosphine), 10 mM CAA (chloroacetamide) and 100 mM Tris-HCl at room temperature. Samples were boiled at 95 °C for 10 min and sonicated for 30 sec for 10 cycles, with 30 sec breaks on high-performance mode with Bioruptor Plus (#B01020001, Diagenode). Samples were centrifuged at 20000 x g for 20 min at RT, supernatants were diluted 10 times with 20 mM Tris and protein concentration was measured using Nanodrop 2000 (#ND-2000, Thermo Fischer Scientific). Three hundred micrograms of each sample were diluted 10 times with 20 mM Tris and digested with 1.5 μl of Mass Spectrometry Grade Trypsin Gold (#V5280, Promega) at 37 °C overnight. Digestion was stopped by adding 50% FA to the reaction at a final concentration of 1%. Samples were centrifuged at 20000 x g for 10 min at RT and supernatants were collected. C-18-SD StageTips were washed and equilibrated sequentially with 200 μl methanol, 200 μl 40% ACN (acetonitrile) / 0.1% FA (formic acid) and 200 μl 0.1% FA by centrifugation, each step for 1 min at RT. Samples were diluted with 0.1% FA, loaded in StageTips and centrifuged for 1-2 min at RT. StageTips were then washed twice with 200 μl 0.1% FA. Tryptic peptides were eluted from StageTips with 100 μl 40% acetonitrile (ACN) / 0.1% formic acid (FA) by centrifugation (300 x g, 4 min, RT). Eluates were dried in a Speed-Vac at 45 °C for 40-45 min and resuspended in 20 μl 0.1% FA. Four micrograms of the peptides were dried in a Speed-Vac and stored at −20°C.

#### TMT10plex labelling

The dried tryptic peptides were reconstituted in 9 μl of 0.1 M TEAB (triethylammonium bicarbonate). Tandem mass tag (TMT10plex; #90110, Thermo Fisher Scientific) labelling was carried out according to the manufacturer’s instructions with the following changes: 0.8 mg of TMT10plex reagent was re-suspended with 70 μl anhydrous ACN. Seven microliters of TMT10plex reagent in ACN were added to 9 μl of clean peptide in 0.1 M TEAB. The final ACN concentration was 43.75% and the ratio of peptides to TMT10plex reagent was 1:20. After 60 min incubation, the reaction was quenched with 2 μl 5% hydroxylamine. Labelled peptides were pooled, dried, re-suspended in 200 μl 0.1% FA, split in two equal parts and desalted using home-made STAGE tips (Li *et al*, 2021).

#### Fractionation of TMT10plex-labeled peptide mixture

One of the two parts was fractionated on a 1 mm x 150 mm ACQUITY column, packed with 130 Å, 1.7 μm C18 particles (#186006935, Waters) using an Ultimate 3000 UHPLC (Thermo Fisher Scientific). Peptides were separated at a flow of 30 μl/min with an 88 min segmented gradient from 1% to 50% buffer B for 85 min and from 50% to 95% buffer B for 3 min; buffer A was 5% ACN, 10 mM ammonium bicarbonate, buffer B was 80% ACN, 10 mM ammonium bicarbonate. Fractions were collected every three minutes, pooled in two passes (fraction 1 + 17, fraction 2 + 18,…, etc.) and dried in a vacuum centrifuge (Eppendorf).

#### LC-MS/MS analysis

Dried fractions were re-suspended in 0.1% FA, separated on a 50 cm, 75 μm Acclaim PepMap column (#164942, Thermo Fisher Scientific) and analyzed on a Orbitrap Lumos Tribrid mass spectrometer (Thermo Fisher Scientific) equipped with a FAIMS device (Thermo Fisher Scientific). The FAIMS device was operated in two compensation voltages, −50 V and −70 V. Synchronous precursor selection based MS3 was used for the acquisition of the TMT10plex reporter ion signals. Peptide separations were performed on an EASY-nLC1200 using a 90 min linear gradient from 6% to 31% buffer; buffer A was 0.1% FA, buffer B was 0.1% FA, 80% ACN. The analytical column was operated at 50 °C. Raw files were split based on the FAIMS compensation voltage using FreeStyle (Thermo Fisher Scientific).

#### Data analysis

Proteomics data was analyzed using MaxQuant, version 1.6.17.0 (Cox & Mann, 2008). The isotope purity correction factors, provided by the manufacturer, were included in the analysis. Differential expression analysis was performed using limma, version 3.34.9 (Ritchie *et al*, 2015) in R, version 3.4.3 (Team, 2017).

### Gene Ontology analysis and Data presentation

Gene Ontology (GO) term enrichment analysis was performed using the Database for Annotation, Visualization and Integrated Discovery (DAVID) tool (Huang da *et al*, 2009a, b) of the Flaski toolbox (Iqbal, 2021) (https://flaski.age.mpg.de, developed and provided by the MPI-AGE Bioinformatics core facility). For the RNA-Seq experiments, either all significantly changing genes (p < 0.05) or selected genes, whose expression levels changed significantly between rapamycin- and DMSO-treated cells with Log_2_-transformed fold change (Log_2_FC) values less than −0.75 (downregulated by rapamycin treatment) or higher than +0.8 (upregulated by rapamycin treatment), roughly corresponding to the top and bottom 5% of the dataset, were used for the DAVID GO analysis (for GOTERM_CC_FAT, GOTERM_BP_FAT). The selection criteria for each analysis are described in the respective figure legends.

For the DAVID GO analysis of quantitative proteomics experiments either all significantly changing proteins (p < 0.05) or selected proteins whose intensity changes significantly between rapamycin- and DMSO-treated cells with Log_2_FC < −0.55 and Log_2_FC > +0.4 (24 h treatments) or Log_2_FC < −0.65 and Log_2_FC > 0.45 (48 h treatments) were used (for GOTERM_CC_FAT, GOTERM_BP_FAT). The human proteome was used as reference list for all analyses.

Cell plots were generated using the DAVID and Cell plot apps in Flaski and include the 15 most significant GO terms from each analysis. The full list of genes/proteins that were detected in each experiment (gray dots) was used for generating the Volcano plots. The respective graphs were prepared using the Scatter plot app in Flaski and labelled in Adobe Photoshop (v. 23.4.2). Significantly-changing genes/proteins are represented by blue (downregulated by rapamycin) or red (upregulated by rapamycin) dots. Proteins/genes whose levels change strongly upon rapamycin treatment (based on the selection criteria described above for each experiment) are shown as blue or red dots with black outline.

## Acknowledgements

We thank all members of the Demetriades lab for critical discussions; Sabine Wilhelm for technical support with sample preparation for proteomics; Ilian Athanasov and Xinping Li from the MPI-AGE Proteomics Core Facility for performing the quantitative proteomics experiments described in this study; the Max Planck Genome Centre (MPGC) Cologne (http://mpgc.mpipz.mpg.de/home/) for performing the RNA-seq experiments described in this study; Franziska Metge and Jorge Boucas from the MPI-AGE Bioinformatics Core for initial RNA-Seq analysis; and the MPI-AGE FACS & Imaging Core Facility for support with cell sorting experiments. FA received support by the Cologne Graduate School of Ageing Research (CGA). CD is funded by the European Research Council (ERC) under the European Union’s Horizon 2020 research and innovation programme (grant agreement No 757729) and by the Max Planck Society. Illustrations in figures created with BioRender.com.

## Author Contributions

Experimental work: FA, NG; data analysis: FA, CD; project design & conceptualization: CD; project supervision: CD; funding acquisition: CD; figure preparation: FA, NG, CD; manuscript draft: FA, CD. All authors approved the final version of the manuscript and agree on the content and conclusions.

## Data availability

The data that support the findings of this study (uncropped immunoblots) or any additional information required to reanalyze the data reported in this paper are available from the corresponding author upon reasonable request. Other datasets that are produced in this study are available in the following databases:

- RNA-Seq data: NCBI Sequence Read Archive (SRA) <u>PRJNA872474</u> (www.ncbi.nlm.nih.gov/sra/PRJNA872474).
- Mass spectrometry proteomics: PRIDE (Perez-Riverol *et al*, 2022) <u>PXD038051</u> (www.ebi.ac.uk/pride/archive/projects/PXD038051).

## Code availability

No code was generated in this study.

## Additional Information

Supplementary Information (Expanded View Figures 1-4 and Supplementary Tables 1-8) is available for this paper.

Correspondence and requests for materials should be addressed to Constantinos Demetriades.

