## Supplementary material for "Comprehensive Evaluation of Rapamycin’s Specificity as an mTOR Inhibitor": Suppl Figures 1-4

### Supplementary Information

Expanded View Figures 1-4 and Supplementary Tables 1-8

### Supplementary Tables

Supplementary Table 1. Differential gene expression analysis in control (WT) and mTOR$^{RR}$ HEK293FT cells upon rapamycin treatment.

Supplementary Table 2. List of genes used for the GO analysis and associated GO terms from the RNA-seq analysis in Rapa- vs DMSO-treated WT cells (only strongly affected genes).

Supplementary Table 3. List of genes used for the GO analysis and associated GO terms from the RNA-seq analysis in Rapa- vs DMSO-treated WT cells (all significantly changing genes).

Supplementary Table 4. Differential protein expression analysis in control (WT) and mTOR$^{RR}$ HEK293FT cells upon rapamycin treatment.

Supplementary Table 5. List of proteins used for the GO analysis and associated GO terms from the proteomic analysis in 24 h Rapa- vs DMSO-treated WT cells (all significantly changing proteins).

Supplementary Table 6. List of proteins used for the GO analysis and associated GO terms from the proteomic analysis in 48 h Rapa- vs DMSO-treated WT cells (all significantly changing proteins).

Supplementary Table 7. List of proteins used for the GO analysis and associated GO terms from the proteomic analysis in 24 h Rapa- vs DMSO-treated WT cells (only strongly affected proteins).

Supplementary Table 8. List of DNA oligonucleotides used in this study.

### Expanded View Figures

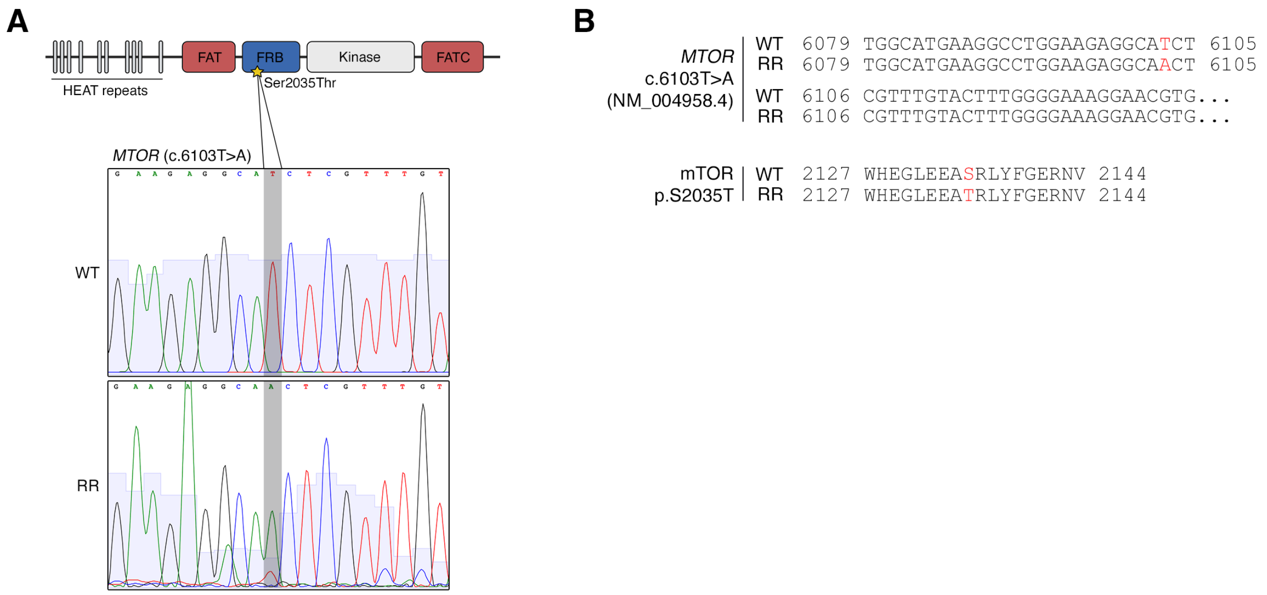


**Figure EV1. Characterization of the *MTOR* genomic alterations in the mTOR$^{RR}$ HEK293FT cells. (A-B)** CRISPR/Cas9-mediated gene-editing of *MTOR*. The associated genomic changes in *MTOR* were validated by Sanger sequencing (A). The resulting changes in the mTOR cDNA and protein sequence are shown in (B). The position of the modified nucleotide / amino acid is shown in red.

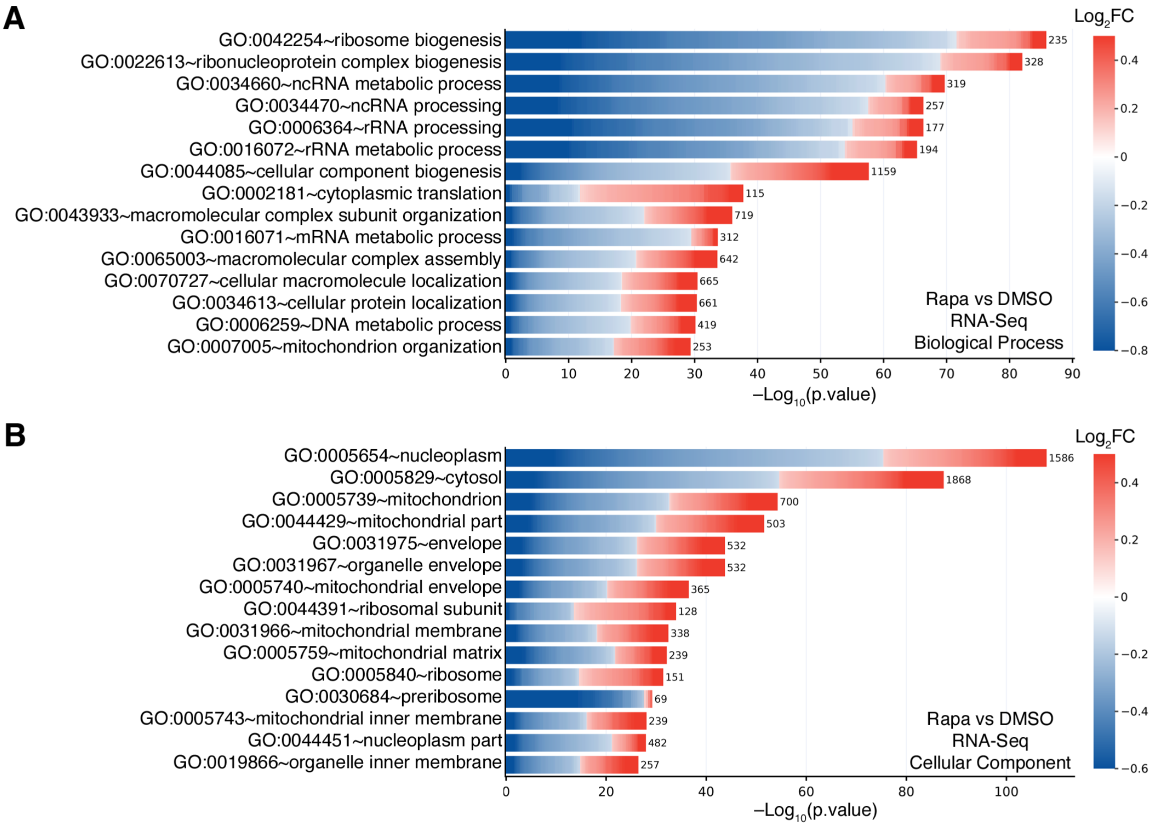


**Figure EV2. GO analysis of the rapamycin-induced changes in the transcriptome of WT HEK293FT cells.**

**(A)** Biological process (BP) GO term analysis using all genes that are significantly ($p < 0.05$) down- (blue) or upregulated (red) by rapamycin in WT cells, as described in Fig. 2B. The color of each box in the cell plot represents log-transformed fold change values for each gene in rapamycin- vs DMSO-treated cells. The number of genes in the selected dataset for each GO term is shown on the right side of each bar.

**(B)** As in (A), but for Cellular Component (CC) GO term analysis.

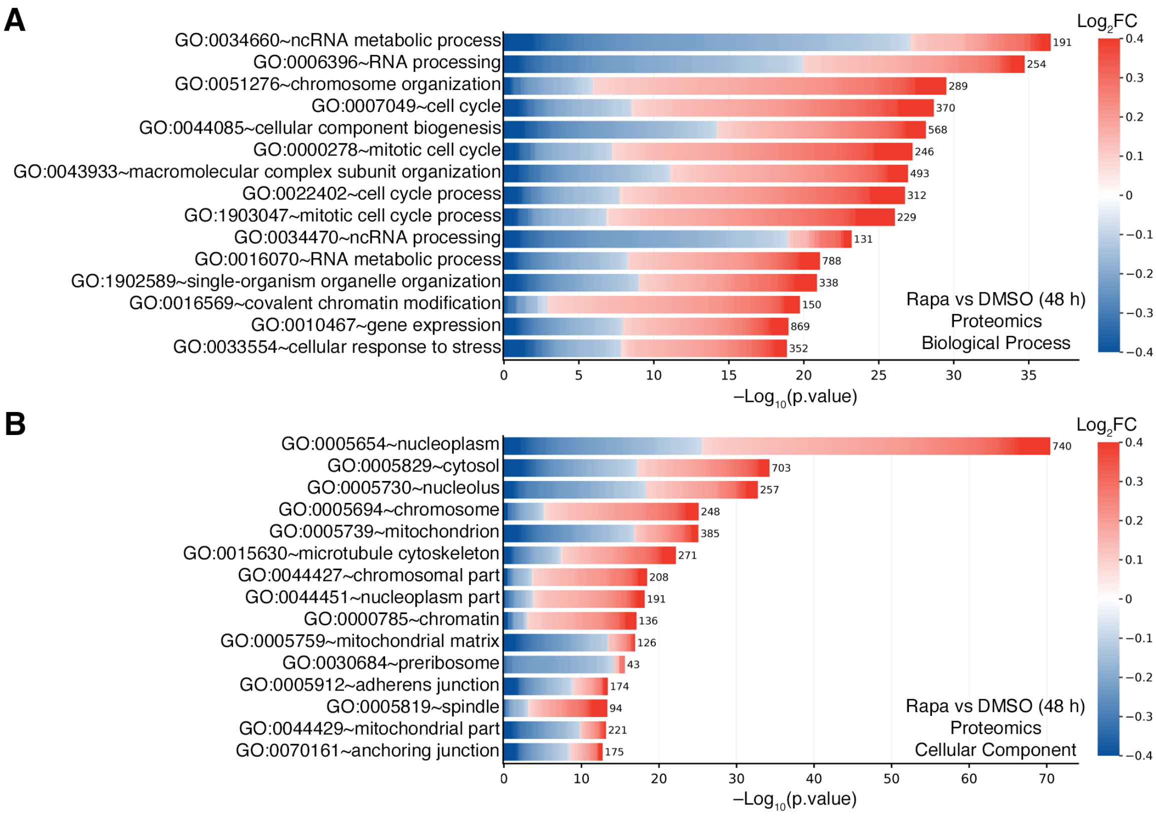


**Figure EV3. GO analysis of the rapamycin-induced changes (48 h) in the proteome of WT HEK293FT cells.**

**(A)** Biological process (BP) GO term analysis using all proteins that are significantly down- (blue) or upregulated (red) by rapamycin (48 h) in WT cells, as described in Fig. 3C. The color of each box in the cell plot represents log-transformed fold change values for each protein in rapamycin- vs DMSO-treated cells. The number of proteins in the selected dataset for each GO term is shown on the right side of each bar.

**(B)** As in (A), but for Cellular Component (CC) GO term analysis.

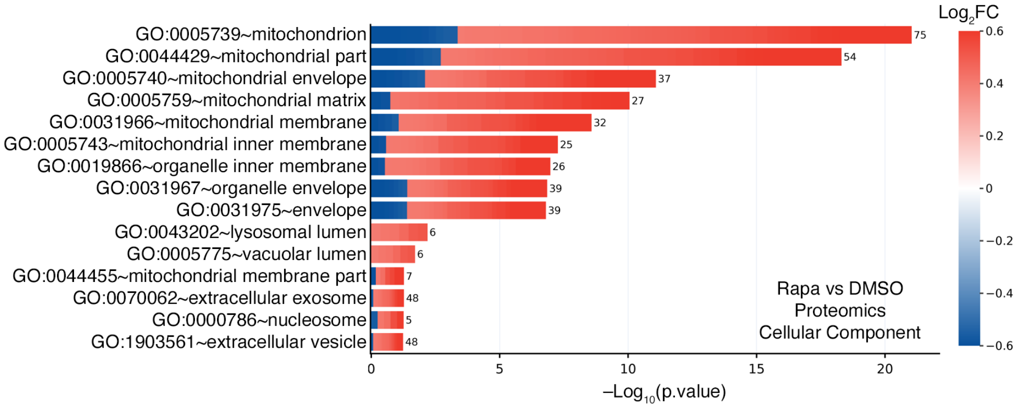


**Figure EV4. GO analysis of the robust rapamycin-induced changes (24 h) in the proteome of WT HEK293FT cells.**

Cellular Component (CC) GO term analysis using only the proteins that are strongly down- (blue) or upregulated (red) by rapamycin (24 h) in WT cells, as described in Fig. 3B. The color of each box in the cell plot represents log-transformed fold change values for each protein in rapamycin- vs DMSO-treated cells. The number of proteins in the selected dataset for each GO term is shown on the right side of each bar.
